# Extracellular plant subtilases dampen cold shock peptide elicitor levels

**DOI:** 10.1101/2024.06.14.599038

**Authors:** Changlong Chen, Pierre Buscaill, Nattapong Sanguankiattichai, Jie Huang, Farnusch Kaschani, Markus Kaiser, Renier A. L. van der Hoorn

## Abstract

Recognizing pathogen-associated molecular patterns (PAMPs) on the cell surface is crucial for plant immunity. The proteinaceous nature of many of these patterns suggests that secreted proteases play important roles in their formation and stability. Here, we demonstrate that the apoplastic subtilase SBT5.2a inactivates the immunogenicity of cold-shock proteins of the bacterial plant pathogen *Pseudomonas syringae* by cleaving within the immunogenic csp22 epitope. Consequently, mutant plants lacking SBT5.2a activity retain higher levels of csp22, leading to enhanced immune responses and reduced pathogen growth. SBT5.2 sensitivity is influenced by sequence variation surrounding the cleavage site and likely extends to CSPs from other bacterial species. These findings suggest that variations in csp22 stability among bacterial pathogens are a crucial factor in plant-bacteria interactions and that pathogens might exploit plant proteases to avoid pattern recognition.

## INTRODUCTION

Pathogen recognition is pivotal to plant survival. Pathogen-associated molecular patterns (PAMPs) are recognized on the plant cell surface by pattern-recognition receptors (PRRs). Upon PAMP binding, PRRs associate with receptor kinases and intracellular protein kinases to activate pattern-triggered immunity (PTI), which includes the production of reactive oxygen species (ROS) and ethylene, apoplast alkalinisation, MAP kinase phosphorylation, and transcriptional reprogramming (Boller and Felix, 2009; Couto and Zipfel, 2016; Jones and Dangl, 2006).

PAMP recognition by PRRs occurs in the extracellular space within plant tissues (the apoplast). The apoplast is the first and often final destination for pathogens and is an important site for pathogen proliferation (Jones & Dangl, 2006). Pathogen colonisation of the apoplast is partly mitigated by plant-secreted hydrolases, including proteases, which are secreted both constitutively and inducibly (Van der Hoorn, 2008; Godson & van der Hoorn, 2021). Ser proteases, including subtilases, are the largest class of secreted proteases (Van der Hoorn, 2008).

PAMPs can be oligosaccharides, lipids and peptides and a wide range of peptide-based PAMPs have been identified from fungal, oomycete and bacterial pathogens (Mott et al., 2014). PRRs tend to perceive a conserved epitope of a PAMP that has important functions to the microbe (Boller & He, 2009). Although the release of immunogenic PAMP peptides from their precursors could be an essential step in pathogen recognition, our knowledge of the biogenesis and maintenance of PAMPs and the involved enzymes is very limited (Chen et al., 2023).

The conserved nucleic acid binding motif RNP-1 of bacterial cold shock proteins (CSPs) serves as a PAMP that is perceived by PRRs from several *Solanaceae* species including tomato (*S. lycopersicum*), tobacco (*N. tabacum*), and potato (*S. tuberosum*) (Felix and Boller, 2003; Saur et al., 2016; Wang et al., 2016). CSPs are highly induced in bacteria in response to rapid downshift in temperature, which is the basis for their name. However, CSPs are also activated by other types of stress and many are constitutively produced (Bae et al., 2000). The 22-residue N-terminal consensus of 150 bacterial CSP sequences from species such as *Micrococcus*, *Bacillus*, *Escherichia* was introduced as csp22. Residues 5-19 of csp22 (csp15) constitute the active PAMP epitope that can trigger plant immune responses, including a burst of reactive oxygen species (ROS) (Felix and Boller, 2003; Saur et al., 2016). Two distinct receptors, the receptor-like protein CSPR and the receptor-like kinase CORE, were claimed to act as CSP receptors in older *N. benthamiana* plants (Saur et al., 2016; Wang et al., 2016), but the affinity of csp22 for CORE is much higher than for CSPR (Wang et al., 2016), and only silencing of *CORE* makes these plants insensitive for csp22 (Dodds et al., 2023). CSPR was later identified as RE02 and is involved in perceiving small Cys-rich proteins from *Valsa mali* (Nie et al., 2020), and *Sclerotinia sclerotiorum* (Yang et al., 2023).

In this study, we tested the hypothesis that extracellular plant proteases might play important roles in the biogenesis of the CSP elicitor of the model bacterial pathogen *Pseudomonas syringae* pv. *tomato* DC3000 (*Pto*DC3000). Surprisingly, however, we discovered proteolytic degradation of csp22 in the apoplast, mediated by serine protease SBT5.2, which dampens the CSP-triggered immune response and suggests that pathogens take advantage of plant proteases to escape PAMP recognition.

## RESULTS

### csp22 from CspD of *Pto*DC3000 is recognized by CORE

To determine the role of CSP elicitor release in bacterial immunity, we studied the release of csp22 from *Pto*DC3000. *Pto*DC3000 has six CSP proteins that contain the cold shock domain (PF00313) and contain peptides similar to the csp22 consensus (**Figure 1A**), of which five are expressed during infection (Nobori et al., 2018) (**Figure 1B**). Since the original discovery of csp22 included the CspD and CapB proteins from various bacteria (Felix and Boller, 2003), we expressed these two proteins in *E. coli* with a His tag linked via the TEV cleavage sequence to the N-terminus. Heterologous expression only delivered sufficient CspD protein (Supplemental **Figure S1**), so we continued our studies with CspD.

**Figure 1.**
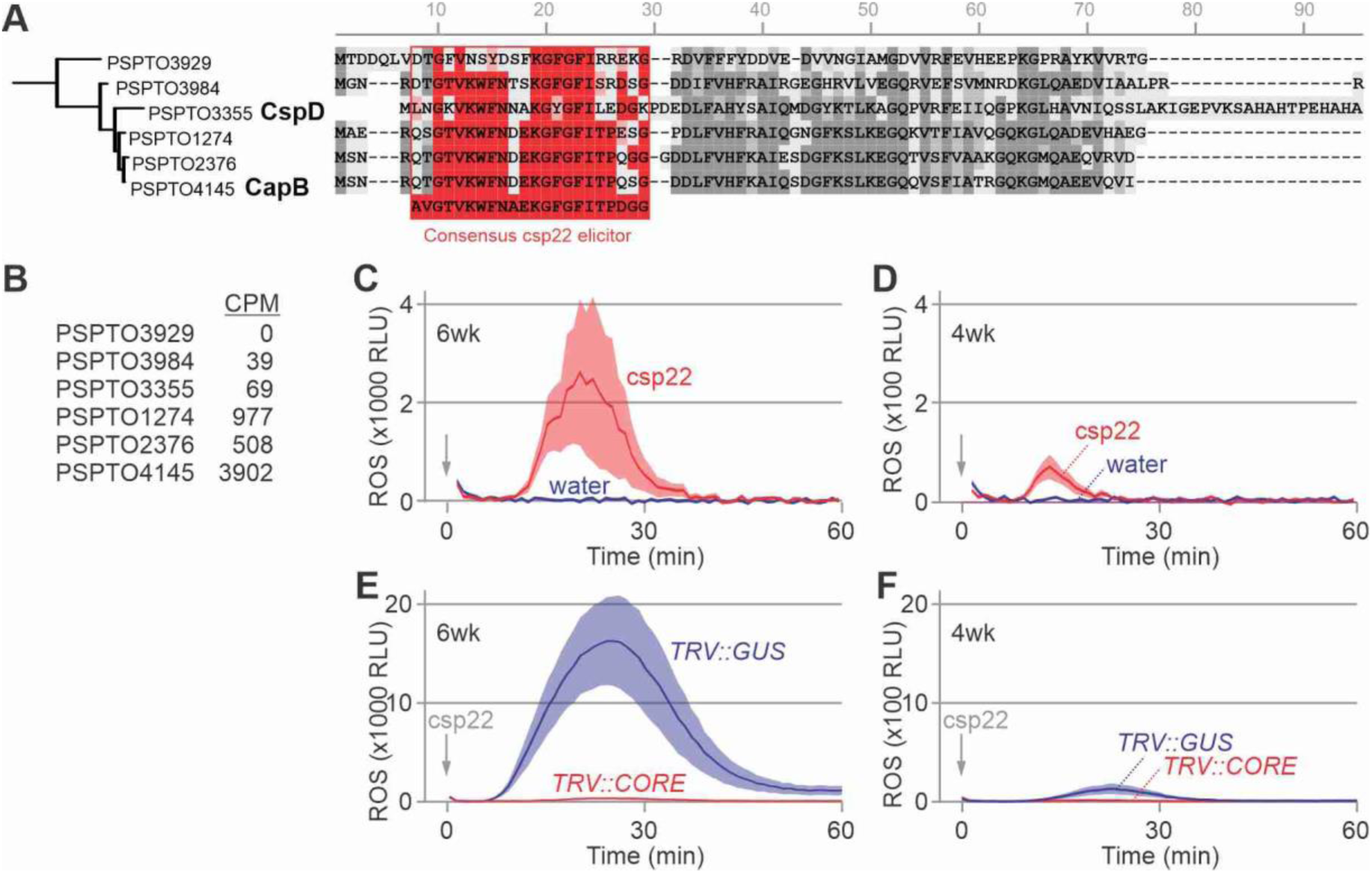
CspD-derived csp22 triggers *Nb*CORE-dependent oxidative burst in *N. benthamiana.* **(A)** Sequence alignment of the six CSPs of *Pto*DC3000 with the frequently used consensus csp22 peptide. **(B)** Transcript levels of CSPs of *Pto*DC3000 during infection in counts per million (CPM). Extracted from (Nobori et al., 2018). **(C)** CspD-derived csp22 triggers an oxidative burst in 6-week old benthamiana plants. **(D)** CspD-derived csp22 triggers only a weak oxidative burst in 4-week old benthamiana plants. **(E)** The oxidative burst triggered by csp22 of CspD in 6wk-old plants is absent upon silencing *Nb*CORE. **(F)** The weak oxidative burst triggered by csp22 of CspD in 4wk-old plants is absent upon silencing *Nb*CORE. (C-F) One-week old *N. benthamiana* plants were infected with and without TRV carrying a fragment of GUS (*TRV::GUS*) or *Nb*CORE (*TRV::CORE*). Leaf discs of 4- and 6-week old plants were treated with water or 500 nM csp22, and ROS was measured in relative light units (RLU). Error shades represent the SE of n=12 (C-D) or n=6 (E-F) leaf discs.

The csp22 peptide of CspD differs at nine residues from the consensus csp22 reported originally (Felix & Boller, 2003). To test if the csp22 peptide of CspD is recognized by *N. benthamiana*, we synthesized this 22-residue peptide and supplemented this to leaf discs of 6wk-old *N. benthamiana* floating on a solution containing luminol and horse radish peroxidase (HRP). Luminescence measurements revealed that csp22 of CspD is able to trigger a classic oxidative burst by releasing reactive oxygen species (ROS), in contrast to the water control (**Figure 1C**). Only a weak oxidative burst was detected in 4wk-old plants (**Figure 1D**), consistent with the low expression of *Nb*CORE in younger plants (Wang et al., 2016).

To confirm that *Nb*CORE is required for detecting csp22, we depleted *Nb*CORE with Virus-induced Gene Silencing (VIGS) using Tobacco Rattle Virus (TRV) vectors carrying a 300bp fragment of *Nb*CORE, or of β-glucuronidase (GUS) as a negative control (Dodds et al., 2023). Oxidative burst assays showed that 6wk-old *TRV::CORE* plants are blind to csp22 of CspD, in contrast to *TRV::GUS* plants, which show an csp22-induced oxidative burst (**Figure 1E**). The weak csp22-induced response detected in 4wk old *TRV::GUS* plants is also absent from *TRV::CORE* plants (**Figure 1F**), indicating that weak responses in younger plants are still *NbCORE*-dependent.

We next tested if also the purified CspD protein (**Figure 2A**) could trigger a *Nb*CORE-dependent oxidative burst. We expressed CspD with a N-terminal His-tag in *E. coli* and purified this over Ni-NTA (**Figure 2A**). Indeed, CspD used at similar concentrations as csp22 triggers an oxidative burst in 6wk-old *TRV::GUS* plants, but not in *TRV::CORE* plants (**Figure 2B**). The oxidative burst was nearly absent from 4wk-old *TRV::CORE* and *TRV::GUS* plants (**Figure 2C**), consistent with the low *Nb*CORE expression in younger plants. These data demonstrate that purified CspD triggers an *Nb*CORE-dependent oxidative burst, indicating that this sample does not contain other elicitors such as the bacterial flagellin of *E. coli*, which could also have triggered an oxidative burst in leaf discs of *N. benthaminana* because they express *Nb*FLS2.

**Figure 2.**
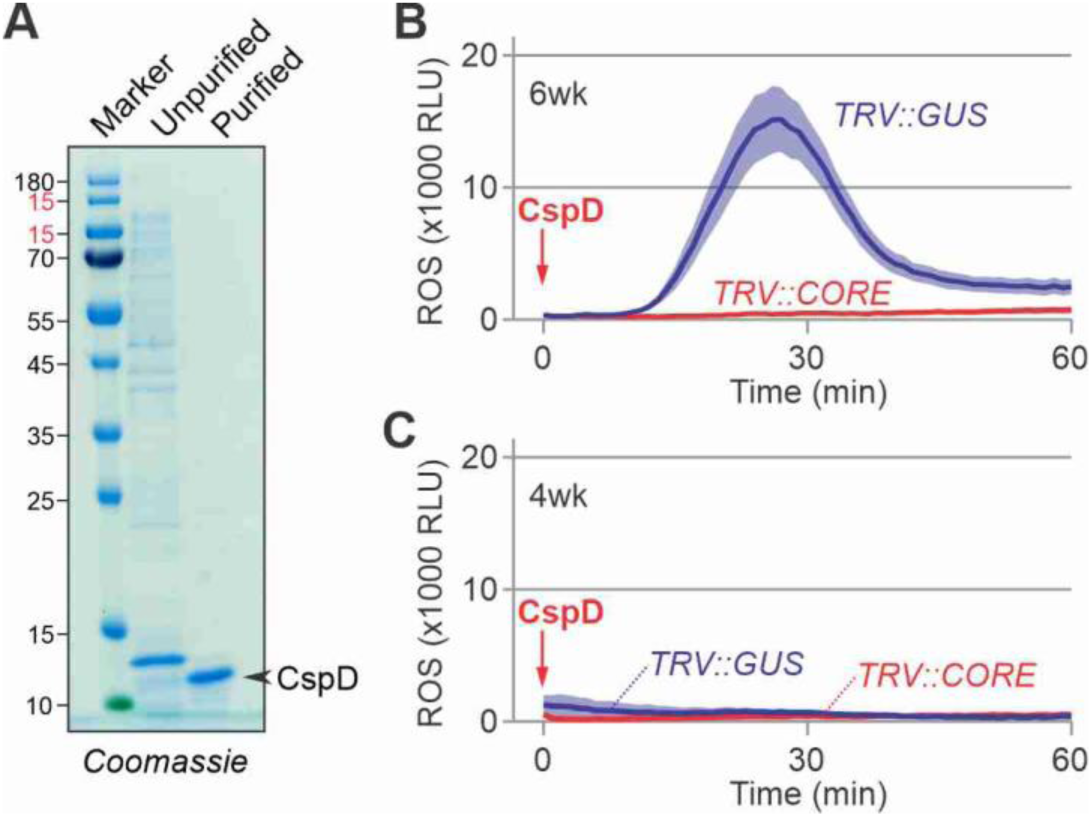
CspD triggers a *Nb*CORE-dependent oxidative burst in 6wk-old *N. benthamiana*. **(A)** Purification of CspD from *P. syringae* heterologously expressed in *E. coli. Pto*CspD was tagged with an N-terminal His-TEV tag, expressed in *E. coli* and purified by immobilised metal affinity chromatography. **(B)** CspD triggers an oxidative burst in 6wk-old plants that is absent upon silencing *Nb*CORE. **(C)** CspD does not trigger an oxidative burst in 4wk-old plants. (B-C) One-week old *N. benthamiana* plants were infected with TRV carrying a fragment of GUS (*TRV::GUS*) or *Nb*CORE (*TRV::CORE*). Leaf discs of 4- and 6-week old plants were treated with 500 nM CspD and ROS was measured in relative light units (RLU). Error shades represent the SE of n = 6 leaf discs.

### Apoplastic fluid quickly degrades CspD

To test whether apoplastic fluids (AFs) could process CspD, we incubated purified CspD with AF isolated from 4∼6wk-old *N. benthamiana* leaves and analysed proteins by SDS-PAGE. This experiment revealed a quick disappearance of intact CspD protein within 15 minutes (**Figure 3A**), indicating that CspD is unstable in AF.

**Figure 3.**
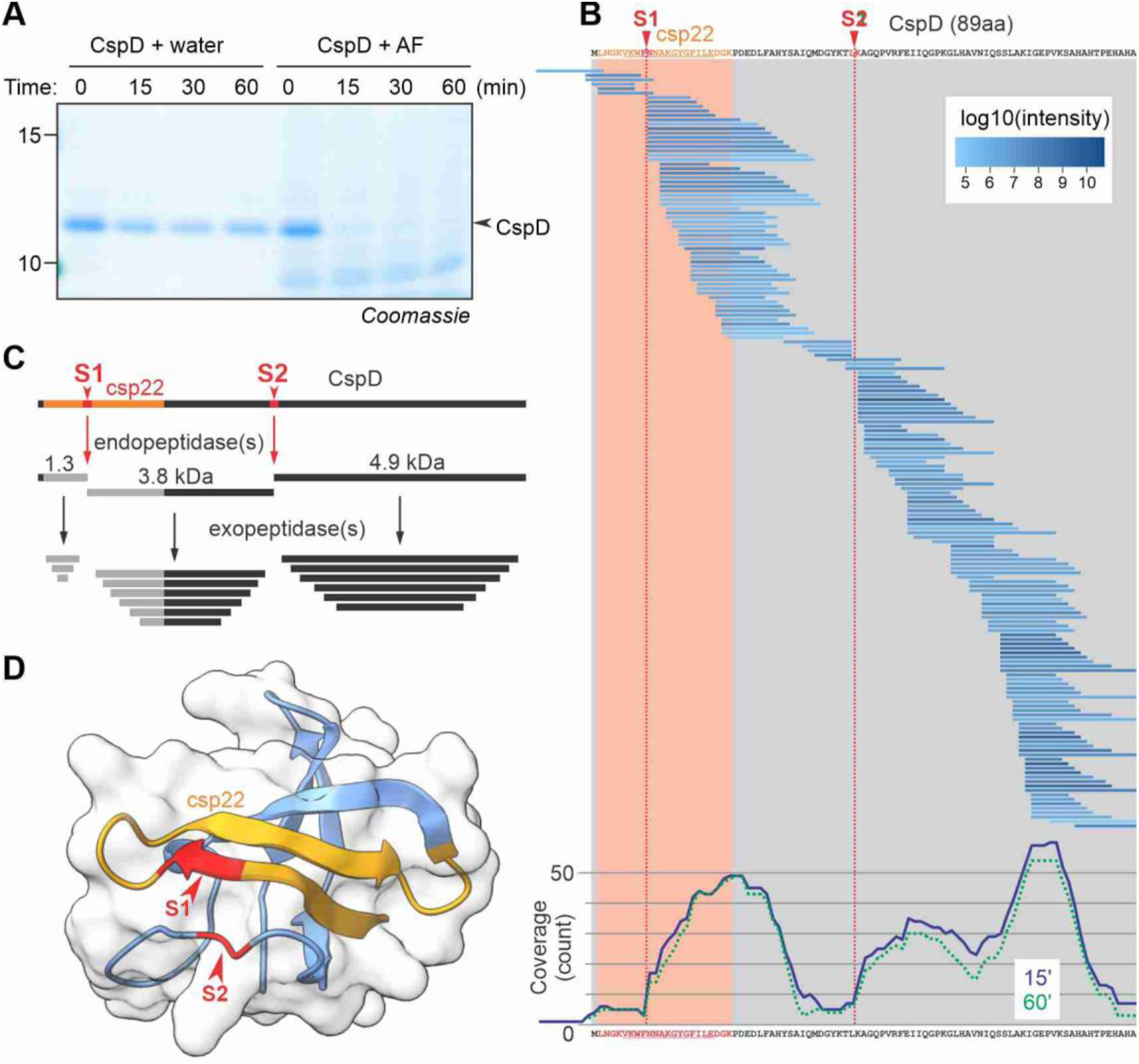
CspD is quickly degraded in apoplastic fluids of *N. benthamiana*. **(A)** Purified CspD is quickly degraded in apoplastic fluid (AF) of *N. benthamiana.* Purified CspD (2 μM) was incubated with water or AF for various times and the products were analyzed on a protein gel by Coomassie staining. **(B)** Degradation products of CspD in AF detected by LC-MS/MS analysis. Purified CspD was incubated with AF isolated from *N. benthamiana* for 15 and 60 minutes. Proteins were precipitated with 80% acetone and the peptide fraction (supernatant) was dried and analyzed by LC-MS/MS. All CspD-derived peptides detected after 15’ incubation were aligned with the CspD protein sequence. The red shaded region indicates csp22 of CspD. Likely initial cleavage sites are indicated by S1 and S2. The bottom graph shows the coverage of the CspD protein after incubation for 15 and 60 (dashed line) minutes, generated by counting the number of times that each residue was detected in the peptides. Peptide coverage graphs of 15 and 60 minutes incubation and the water controls are shown in Supplemental **Figure S2. (C)** Proposed degradation of CspD, first by endopeptidase(s) at sites S1 and S2 and then by exopeptidases. Cleavage at S1 inactivates the csp22 elicitor peptide. **(D)** position of the S1 and S2 sites in the structural model of CspD. This structural model was obtained from the Alphafold-predicted structure of CspD in the Uniprot database (Entry Q87ZR9), trimmed for the cold shock domain, with the csp22 peptide (orange) and S1 and S2 cleavage sites (red) as indicated.

To investigate CspD degradation in AF further, we analysed the released peptides by LC-MS/MS and mapped the CspD-derived peptides onto the CspD protein sequence. A total of 171 different CspD-derived peptides were detected, covering the entire CspD protein sequence (**Figure 3B**). Many peptides overlap and are staggered in three clusters, differing only in single residues removed from the N- and C-terminus (**Figure 3B** and Supplemental **Figure S2**). This peptide pattern indicates that CspD is cleaved at two sites (S1 and S2) by endopeptidases cleaving at VKWF↓NNAK and YKTL↓KAGQ, and that exopeptidases subsequently remove a series of single residues from either end (**Figure 3C**). Cleavage at site S1 would cleave 10.0 kDa CspD into fragments of 1.3 and 8.7 kDa, whereas processing at S2 would result in 5.1 and 4.9 kDa fragments (**Figure 3C**). Analysis of the CspD structural model (Chen et al., 2023) indicates that site S1 locates in the exposed β-sheet in middle of the csp22 elicitor sequence, and that site S2 locates in a nearby exposed loop (**Figure 3D**).

### AF quickly inactivates csp22

Cleavage at site S1 of CSP within the csp22 epitope was unexpected and would inactivate the immunogenicity of csp22. To confirm csp22 cleavage, we incubated the csp22 peptide briefly in AF and then added this to leaf discs of csp22-responsive plants to detect remaining elicitor activity. Remarkably, elicitor activity drastically diminished within five minutes and was even absent after 15 minutes incubation (**Figure 4A**), demonstrating that csp22 is unstable in AF.

**Figure 4.**
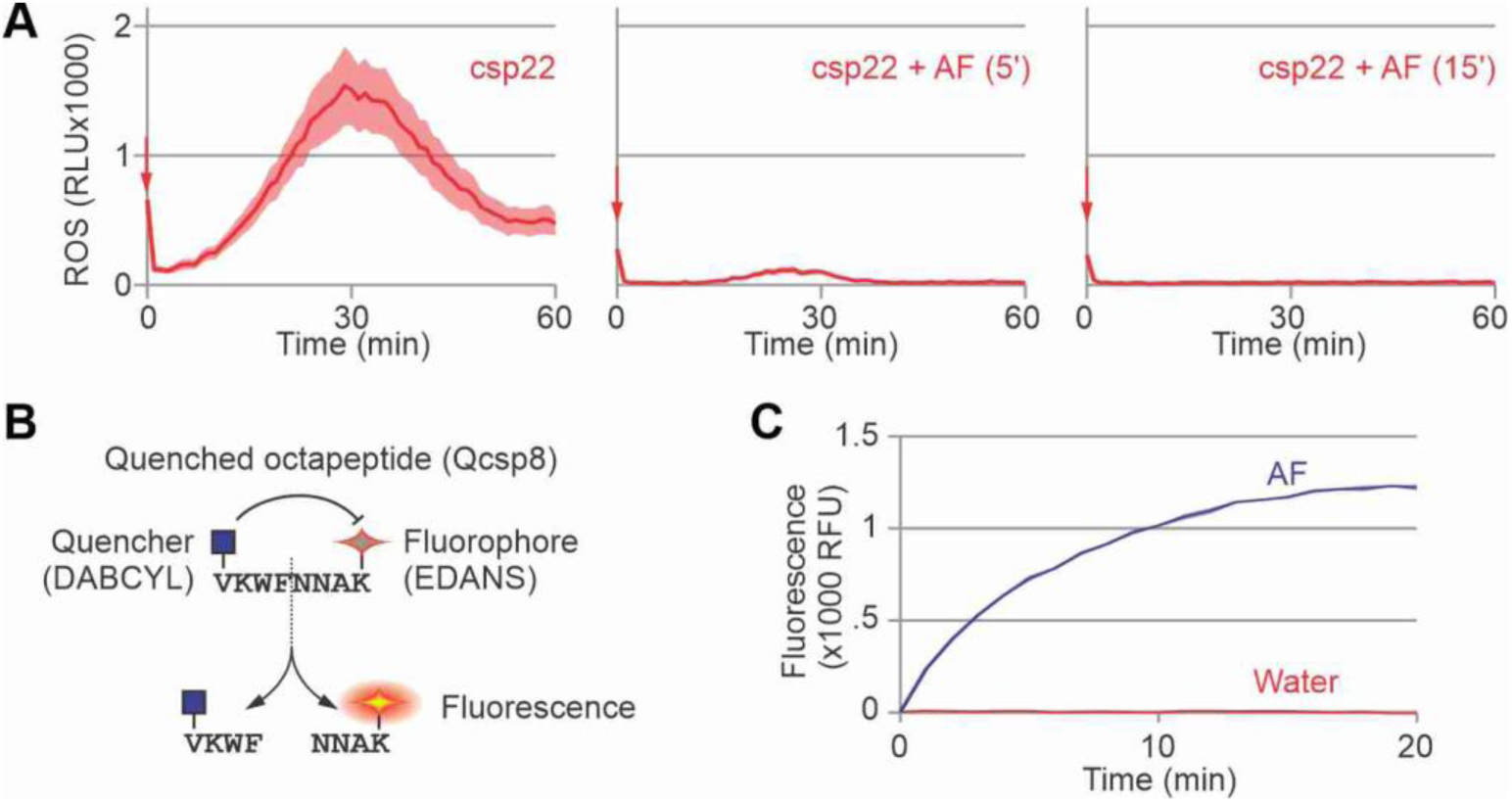
csp22 is quickly inactivated in apoplastic fluids. **(A)** Incubation in AF quickly inactivates the csp22 elicitor. AF was incubated with 500 nM csp22 peptide for 5 or 15 minutes and then added to leaf discs of 6wk-old *N. benthamiana* plants floating on the luminol-HRP solution. Luminescence (relative light unites, RLU), was measured for 60 minutes with a platereader. Error shades represent SE of n=12 replicates. **(B)** Concept of quenched csp8 octapeptide reporting the cleavage in csp22 peptide. **(C)** Qcsp8 is quickly cleaved in AF isolated from *N. benthamiana* leaves. Qcsp8 (10 μM) was incubated in water or AF and fluorescence was measured over time. Error shades represent SE of n = 3 replicates.

To confirm cleavage in the middle of csp22, we obtained a custom-synthesised quenched octapeptide containing the eight residues surrounding the S1 site (VKWF↓NNAK). Cleavage of this Qcsp8 peptide would separate the N-terminal quencher (DABCYL) from the C-terminal fluorophore (EDANS), causing fluorescence (**Figure 4B**). Incubation of Qcsp8 in AF causes a rapid increase of fluorescence compared to the water control (**Figure 4C**), indicating that this octapeptide is cleaved in AF.

### Subtilases inactivate csp22, degrade CspD and cleave Qcsp8

Being the largest class of proteases detected in the apoplast of *N. benthamiana* (Grosse-Holz et al., 2018), subtilases were tested for degrading csp22/CspD/Qcsp8 by taking advantage of subtilase inhibitor Epi1 of the oomycete potato blight pathogen *Phytophthora infestans* (Tian et al., 2004). Elicitor activity was detected after incubation of csp22 peptide with AFs isolated from leaves transiently expressing Epi1 (AF(Epi1)), but not from leaves transiently transformed with the Empty Vector control (AF(EV)) (**Figure 5A**), indicating that Epi1 blocks AF degradation of csp22, implying that apoplastic subtilases are responsible for degrading csp22.

**Figure 5.**
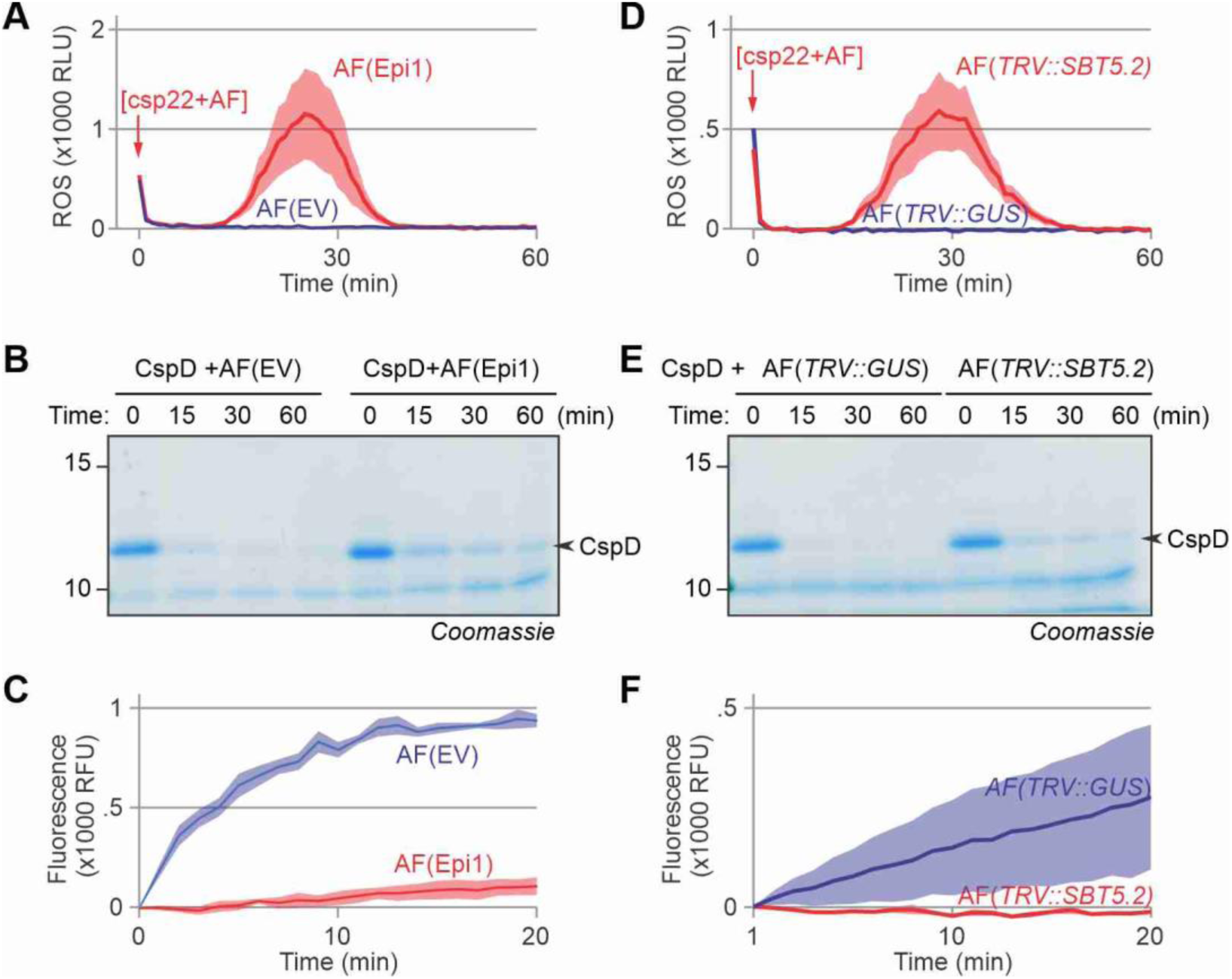
SBT5.2 subtilases are required for inactivating csp22, degrading CspD and cleaving Qcsp8. **(A)** Inactivation of csp22 is suppressed in AF of plants overexpressing subtilase inhibitor Epi1. **(B)** CspD degradation is reduced in AF of plants overexpressing subtilase inhibitor Epi1. **(C)** Cleavage of Qcsp8 is reduced in AF of plants overexpressing subtilase inhibitor Epi1. **(D)** Inactivation of csp22 is suppressed in AF isolated from *TRV::SBT5.2* plants. **(E)** CspD degradation is reduced in AF isolated from *TRV::SBT5.2* plants. **(F)** Cleavage of Qcsp8 is reduced in AF isolated from *TRV::SBT5.2* plants. **(A,D)** AF was incubated with 500 nM csp22 peptide for 60 minutes and then added to leaf discs of 6wk-old *N. benthamiana* plants floating on a luminol-HRP solution. Luminescence (relative light units, RLU), was measured for 60 minutes with a plate reader. Error shades represent SE of n=6 and 8 replicates for (A) and (D), respectively. **(B,E)** AF was incubated with 2 μM CspD protein for various time points and separated on a 15% SDS PAGE gel and stained with Coomassie. **(C,F)** AF was incubated with 10 µM Qcsp8 whilst fluorescence was measured with a plate reader. Error shades represent SE of n=4 samples.

We also tested if CspD protein is more stable in the presence of Epi1 and found that CspD protein is stabilised in AF containing Epi1 and still degraded in AF of the EV control (**Figure 5B**). Likewise, AF containing Epi1 was no longer able to cleave Qcsp8 when compared to the AF(EV) control (**Figure 5C**), indicating that subtilases are responsible for Qcsp8 cleavage. Taken together, these data indicate that subtilases are responsible for inactivating csp22, degrading CspD protein and cleaving Qcsp8.

### Subtilase SBT5.2 is required for inactivating csp22, degrading CspD and cleaving Qcsp8

Being the most active subtilase in the apoplast of *N. benthamiana* (Grosse-Holz et al., 2018; Jutras et al., 2019; Puchol Tarazona et al., 2021; Paulus et al., 2020; Sueldo et al., 2024; Beritza et al., 2024), subtilase SBT5.2 was further investigated for its role in csp22 degradation. We used VIGS to suppress *Nb*SBT5.2 expression in *N. benthamiana* (Paulus et al., 2020; Beritza et al., 2024) and extracted AF from *TRV::SBT5.2* plants. *N. benthamiana* expresses three SBT5.2 homologs (a-c), that are all targeted with the 300bp fragment of *SBT5.2a* present in *TRV::SBT5.2* (Beritza et al., 2024). Elicitor activity of csp22 was detected upon incubation in AF from *TRV::SBT5.2* plants, but not in AF of *TRV::GUS* control plants (**Figure 5D**), indicating that SBT5.2 is required for inactivating csp22. Likewise, CspD protein was more stable in AF of *TRV::SBT5.2* plants when compared to AF of *TRV::GUS* control (**Figure 5E**), and the Qcsp8 peptide is no longer cleaved in AF of *TRV::SBT5.2* plants, when compared to the *TRV::GUS* control (**Figure 5F**). Taken together, these data indicate that apoplastic subtilase SBT5.2 is required for inactivating csp22, degrading CspD protein and cleaving Qcsp8.

### Purified SBT5.2a inactivates csp22, processes CspD and cleaves Qcsp8

To determine if SBT5.2 is also sufficient for inactivating csp22, degrading CspD protein and cleaving Qcsp8, we cloned the open reading frame encoding SBT5.2a (NbD038072 in NbDE database (Kourelis et al., 2019); NbL13g04590 in LAB360 database (Ranawaka et al., 2023)) fused to a C-terminal His tag into a binary vector and purified this protein from the AF of agroinfiltrated plants on Ni-NTA columns (Supplemental **Figure S3**), resulting in purified SBT5.2a-His protein (**Figure 6A**). We detected multiple isoforms that are labelled with the activity-based Ser hydrolase probe FP-TAMRA (**Figure 6A**), indicating that they are derived from active proteases. Incubation of csp22 with different concentrations of purified SBT5.2a-His resulted in a dose-dependent inactivation of csp22 elicitor activity (**Figure 6B**). Likewise, incubation of CspD with different concentrations of purified SBT5.2a-His resulted in a dose-dependent cleavage of CspD into a smaller isoform that was probably caused by cleavage at the S1 site because that cleavage would result in a 8.7 kDa product, whereas processing at S2 would generate two ∼5 kDa products (**Figure 6B**). Finally, incubation of Qcsp8 with different concentrations of purified SBT5.2a-His resulted in a dose-dependent cleavage of Qcsp8 (**Figure 6D**). By contrast, Qcsp8 was not cleaved by purified P69B-His (**Figure 6E**), a subtilase from tomato (Paulus et al., 2020), demonstrating the specificity of Qcsp8 cleavage by SBT5.2a. Taken together, these data indicate that apoplastic subtilase SBT5.2a can inactivate csp22, process CspD protein and cleave Qcsp8.

**Figure 6.**
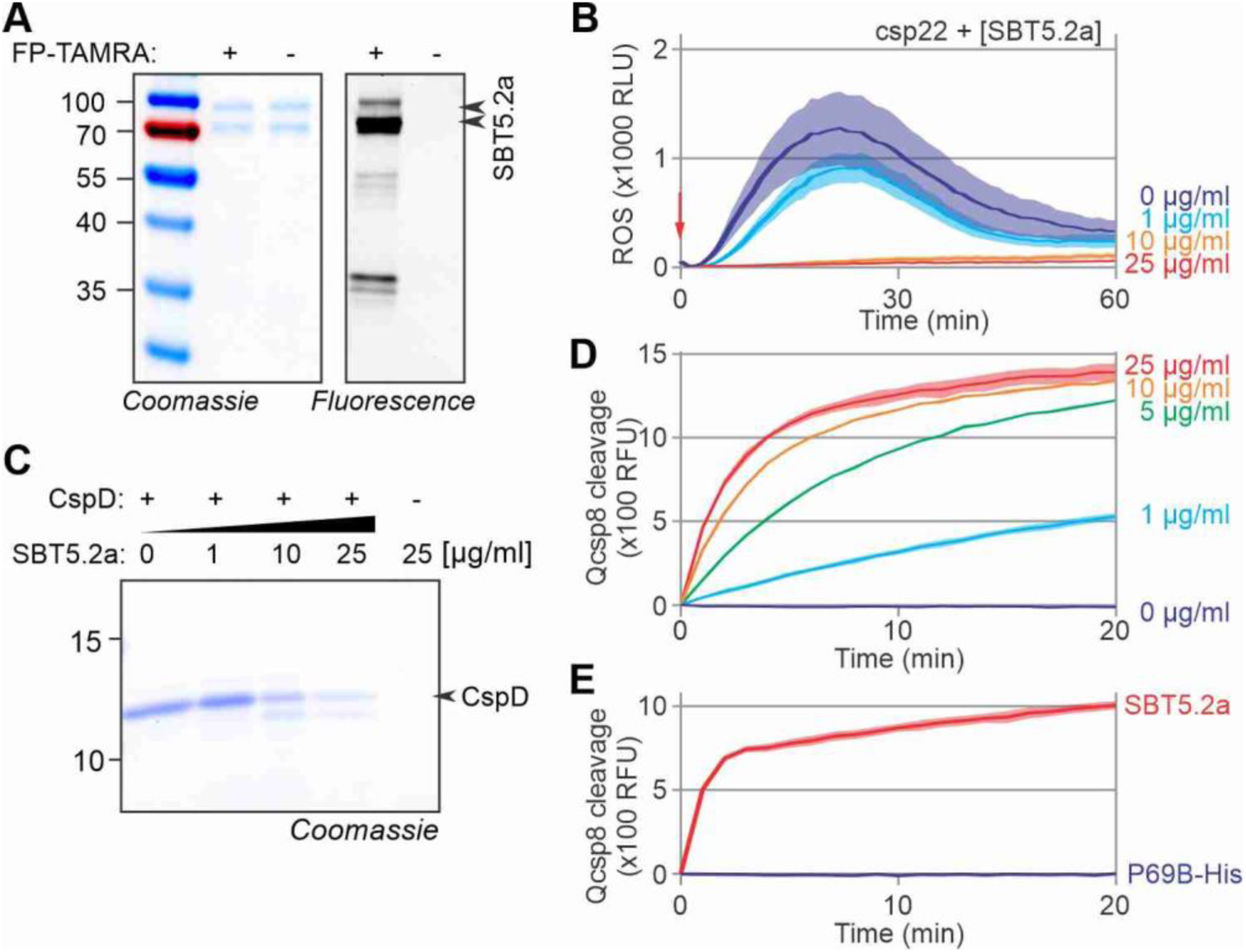
Purified SBT5.2a inactivates csp22 and cleaves CspD and Qcsp8. **(A)** Purified SBT5.2a-His. His-tagged SBT5.2a was transiently expressed in *N. benthamiana* by agroinfiltration and isolated from AF at day six on Ni-NTA columns. Proteins were labeled with and without 0.2 μM FP-TAMRA for one hour and separated on SDS-PAGE stained with Coomassie or scanned for in-gel fluorescence. **(B)** Purified SBT5.2a-His inactivates the csp22 elicitor. Purified SBT5.2a-His was incubated with 100 nM csp22 peptide for 45 minutes and then added to leaf discs of 6wk-old *N. benthamiana* plants floating on a luminol-HRP solution. Luminescence (relative light units, RLU), was measured for 60 minutes with a plate reader. Error shades represent SE of n=6 replicates. **(C)** Purified SBT5.2a-His processes purified CspD protein. Purified CspD protein was incubated with various concentrations of purified SBT5.2a-his at room temperature for 40 minutes and separated on SDS-PAGE and stained with Coomassie. **(D)** SBT5.2a cleaves Qcsp8. Various concentrations of purified SBT5.2a-His were incubated with 10 μM Qcsp8 whilst fluorescence was measured using a plate reader. Error shades represent SE of n=3 replicates. **(E)** Qcsp8 is cleaved by SBT5.2a-His but not by P69B-His. 0.5 μg of purified SBT5.2a-His and P69B-His were incubated with 10 μM Qcsp8 whilst fluorescence was measured using a plate reader. Error shades represent SE of n=4 replicates.

### Mutant *sbt5.2* plants have enhanced immune priming capacity

To investigate the role of SBT5.2 subtilases in immunity, we took advantage of our recently established triple knockout line lacking all three *SBT5.2* genes generated by genome editing (Beritza et al., 2024). AF from these mutants are unable to cleave Qcsp8 (**Figure 7A**) nor inactivate csp22 (**Figure 7B**). To assess the role of SBT5.2 in csp22-mediated immunity we used the *fliC* mutant of *Pto*DC3000*ΔhopQ* (*Pto*DC3000*ΔhopQΔfliC*, Kvitko et al., 2009) to avoid responses triggered by flagellin/FLS2 signalling.

**Figure 7.**
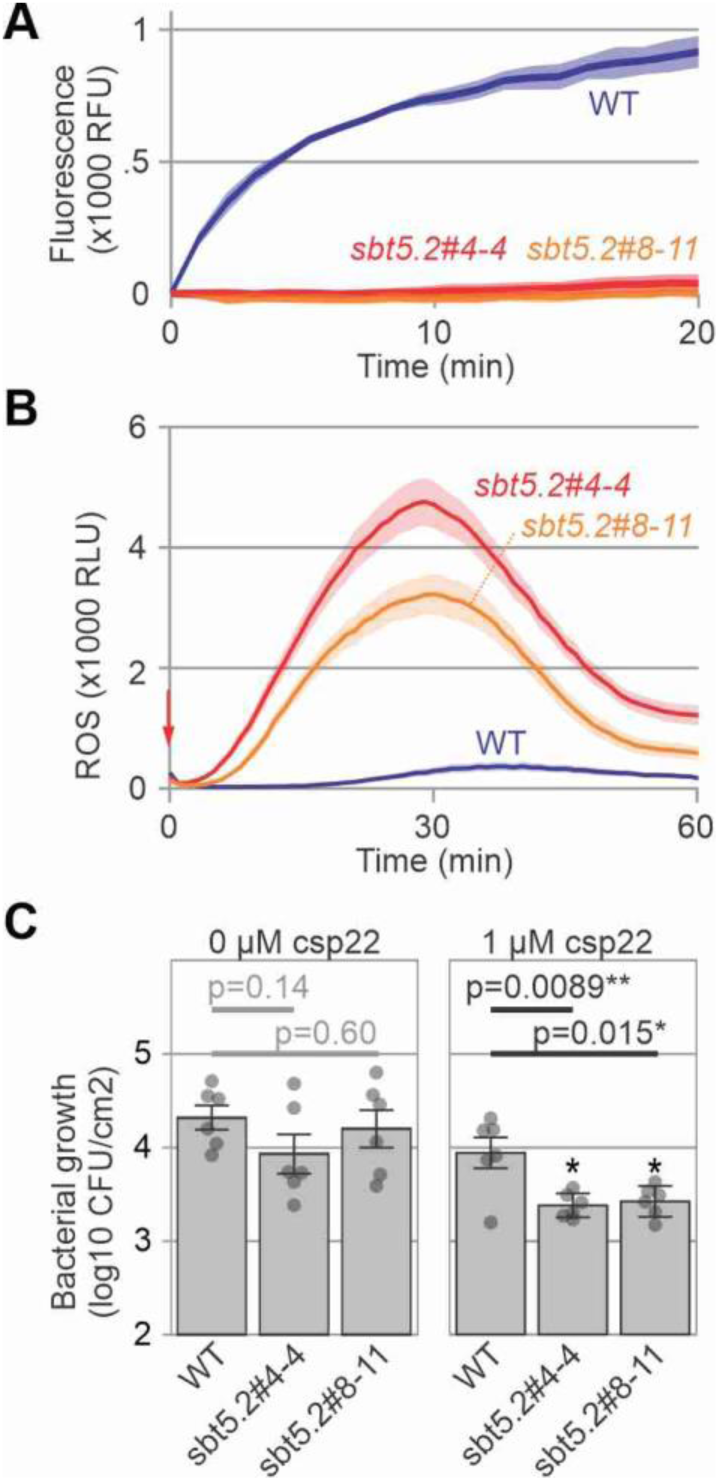
Increased stability of csp22 in *sbt5.2* triple mutants increases immunity. **(A)** The quenched csp8 octapeptide is no longer cleaved in AF of *sbt5.2* mutants. Quenched octapeptide (10 µM) was incubated with AF of WT and *sbt5.2* mutant plants and fluorescence was measured with a plate reader. Error shades represent SE of n=4 samples. **(B)** Csp22 inactivation is reduced in AF isolated from both *sbt5.2* triple mutants. AF of WT or *sbt5.2* mutant plants were incubated with 500 nM csp22 peptide for 30 minutes then added to leaf discs of 6wk-old *N. benthamiana* plants floating on the luminol-HRP solution. Luminescence (relative light unites, RLU), was measured for 60 minutes with a plate reader. Error shades represent SE of n=12 replicates. **(C)** Immunity induced by low csp22 concentration is increased in the *sbt5.2* mutant. Leaves of WT and *sbt5.2* mutant *N. benthamiana* were infiltrated with water or 1 μM csp22 and incubated for 24 hours and then injected with 1x10^5^ bacteria/mL of the flagellin mutant strain of *Pseudomonas syringae* pv. *tomato* DC3000 (*Pta* DC3000*ΔfliCΔhopQ*). Colony forming units (CFUs) were determined one day post infection (dpi). Error bars represent SE of n=6 replicates. P-values are from an unpaired, two-tailed Students *t*-test.

Infection assays, however, did not reveal an altered bacterial growth on the *sbt5.2* mutant lines compared to WT plants (**Figure 7C**). Elicitation with csp22 did also not reveal any changes in the oxidative burst (Supplemental **Figure S4A**), even at low csp22 concentrations (Supplemental **Figure S4B**). We then tested if immune priming (Zipfel et al., 2004) by csp22 is enhanced in *sbt5.2* mutants. Pre-treatment with 1µM csp22, followed by pathogen inoculation after 24 hrs reduces bacterial growth in both *sbt5.2* mutant lines compared to WT plants (**Figure 7C**). Likewise, pretreatment with 5 μM csp22 also reduces bacterial growth of the *fliC* mutant of *Pta*6605 (*Pta*6605*ΔfliC*, Shimizu et al., 2003) more in *sbt5.2* mutant plants when compared to WT plants (Supplemental **Figure S5**). These data demonstrate that the degradation of csp22 by SBT5.2 in WT plants increases plant susceptibility to bacterial infection.

### Polymorphisms in csp22 dictate SBT5.2 sensitivity

To investigate if other csp22 elicitor peptides present in CSP proteins of *Pto*DC3000 have differential sensitivity to SBT5.2-mediated inactivation, we tested two more csp22 peptides that are distinct from csp22 of CspD: csp22 of CapB (PSPTO4145), and PSPTO3984, which differs 11 and 10 residues from csp22 of CspD, respectively (**Figure 8A**). Both csp22 peptides trigger ROS bursts in leaf discs of 6wk-old *N. benthamiana* plants (Supplemental **Figure S6**). All three peptides are inactivated when incubated in AF, but the speed of inactivation differs per peptide (**Figure 8B**). When incubated with AF of WT plants, the csp22 peptide of PSPTO3984 is more quickly degraded than that of CspD, whereas the degradation of csp22 of CapB is slower than CspD (**Figure 8B**). However, all peptides are stabilised to similar levels when incubated in AF of the *sbt5.2* mutant (**Figure 8B**), indicating that SBT5.2 subtilases contribute to the degradation of all three csp22 peptides. Because the residues preceding the cleavage site are identical between CspD, PSPTO3984 and CapB, the reduced processing of CapB by SBT5.2 is probably caused by the presence of acidic residues (D16 and E17) after the cleavage site. These data indicate that the polymorphism in csp22 peptides can lead to varying sensitivity to SBT5.2-mediated cleavage, resulting in different stabilities in the plant apoplast. The alignment with csp22 peptides from different bacterial plant pathogens indicates that the variation in csp22 stabilities might be similar between different plant species (**Figure 8C**).

**Figure 8.**
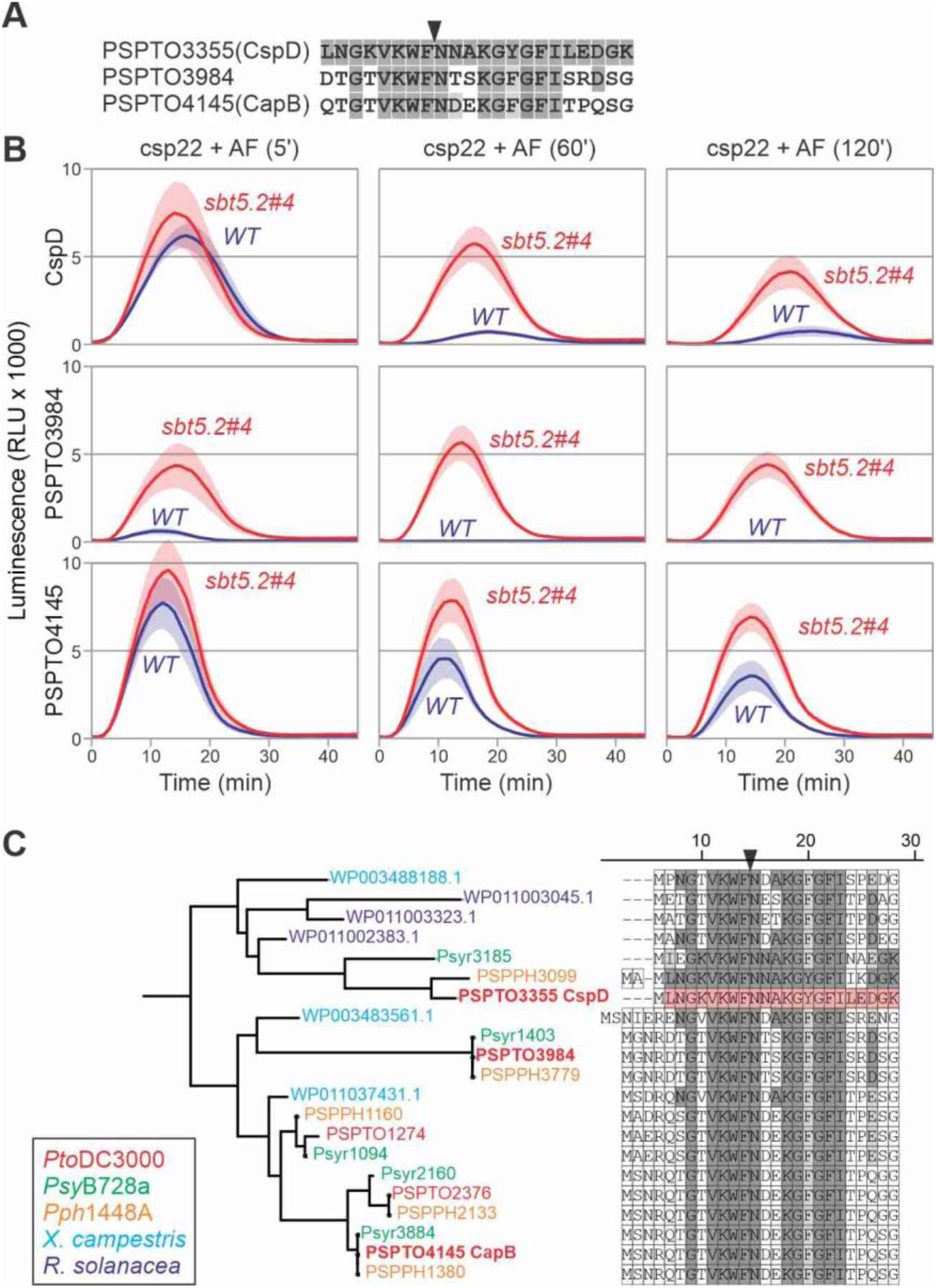
Differential processing of csp22 peptides by SBT5.2. **(A)** Alignment of csp22 sequences from CspD (PSPTO03355), PSPTO3984 and PSPTO4145 from *Pto*DC3000. **(B)** The csp22 peptides (1,000 nM) were incubated with AF from WT plants and the *sbt5.2#4* mutant for 5, 60 and 120 min, 10-fold diluted, and then added to leaf discs of *N. benthamiana* floating in HRP and luminol. ROS burst was measured immediately with a plate reader over 45 min. Error shades indicate SE of n=8 replicates. **(C)** Phylogeny of CSPs with an alignment of enclosed csp22 sequences of *Pseudomonas syringae* pv. *tomato* DC3000 (*Pto*DC3000, red), pv. *syringae* B728a (*Psy*B728a, green) and pv. *phaseolicola* 1448A (*Pph*1448a, orange), as well as *Xanthomonas campestris* ATCC33913 (cyan) and *Ralstonia solanacea* GMI1000 (blue). Highlighted residues are identical (dark grey) or similar (light grey) to the csp22 sequence of CspD.

## DISCUSSION

This work demonstrated that immunogenic csp22 peptides are quickly inactivated by subtilase SBT5.2 in the apoplast. This discovery indicates that SBT5.2 dampens PTI by degrading csp22 and that bacteria may take advantage of SBT5.2 degrading elicitors to evade recognition.

This project was initiated to investigate if peptide elicitors require proteolytic processing for their release from their precursors. We anticipated this since most peptide elicitors are not accessible within the folded precursor (Chen et al. 2023), and many peptide hormones have to be released from precursors to be perceived elsewhere by receptors (Stintzi & Schaller, 2022; Yang et al., 2023). The CSP protein, however, is very small and much of the csp22 peptide is already exposed (**Figure 3D**). In addition, since the csp22 peptide is at the N-terminus, only C-terminal processing would be required for csp22 release. However, the timing of the oxidative burst upon CspD is not significantly different from that of csp22 (**Figure 1CE** and 2B) and we were unable to block CspD perception with subtilase inhibitors PMSF or Epi1 (Supplemental **Figure S7**). And although csp22 binding to CORE has been demonstrated (Wang et al., 2016), unprocessed CSP proteins are likely to bind to CORE, similar to how flagellin protein can bind FLS2 (Meindl et al., 2000). The hypothesis that processing is not required for CSP perception is also consistent with the distribution of detected peptides over the CspD protein when incubated with AF as the detected peptide pattern suggests that there were only two initial processing sites (S1 and S2, **Figure 3B**). Whereas processing as site S1 inactivates csp22, site S2 is far beyond the csp22 peptide. Furthermore, incubation of purified CspD with purified SBT5.2a indicates that site S1 is cleaved first and cleavage at S2 might be a secondary cleavage event (**Figure 6C**). These observations indicate that processing might not be important for the release of the csp22 elicitor. By contrast, the inactivation of the csp22 epitope by SBT5.2 at site S1 seemed a more relevant process.

Indeed, subsequent experiments with csp22 confirmed that csp22 is unstable in the apoplast. We provided multiple lines of evidence that SBT5.2s are both required and sufficient for inactivating csp22. Inactivation of csp22 in AF can be blocked with SBT-inhibitor Epi1, by silencing *SBT5.2s*, or by inactivating *SBT5.2s* by genome editing. Conversely, purified SBT5.2a is sufficient to inactivate csp22 and can cleave the quenched octapeptide carrying the presumed S1 cleavage site. Also the processing of CspD protein by purified SBT5.2a suggests that the S1 site is cleaved by SBT5.2. The S1 site in csp22 (VKWF↓NNAK) is consistent with the known promiscuity of this protease. SBT5.2 is also required for cleaving the prodomain of proRcr3 in the region carrying the sequence (EFKI-)NDLSDDYM(-PSN) irrespective of random mutagenesis in this region (Paulus et al., 2020), and purified SBT5.2a was found to cleave various recombinant proteins at sites TTLF↓GVPI, YNYY↓DFYD, YNYH↓YMDV, YHYM↓DVWG, TTVT↓VSSA, VTVS↓SAST, MFLE↓AIPM and AIPM↓SIPP (Puchol Tarazona et al., 2021). Although there are often hydrophobic residues at positions P4, P2 and P1 (V, W and F in csp22, respectively), there are several exceptions, indicating that SBT5.2 might be promiscuous and it is rather the combination of residues in a peptide that may dictate SBT5.2 cleavability. The promiscuity of SBT5.2 is consistent with observations made for Arabidopsis SBT5.2, which was found to process the precursors of Inflorescence Deficient in Abscission (proIDA, Schardon et al., 206), and Epidermal Patterning Factor (proEPF2, Engineer et al., 2014), as well as flagellin (Matsui et al., 2024), at sites that share not much homology.

The consequence of csp22 processing is that the csp22 levels are suppressed by SBT5.2 and that, consequently, the immune response is dampened by SBT5.2s. Indeed, priming by low csp22 concentrations induces PTI only in *sbt5.2* mutants (**Figure 7C**), confirming that SBT5.2 dampens csp22-triggered PTI. A dampened immune response may benefit the plant by restricting costly immune responses to the site of pathogen infection by avoiding the diffusion of peptide elicitors. Likewise, csp22 degradation will also result in a more temporal immune response, so that PTI is no longer triggered in plants after an infection has cleared. Both temporal and spatial dampening of immune response would avoid unnecessary activation of costly immune responses and make more resources available to promote plant growth.

But that SBT5.2s inactivate csp22 is also a clear benefit to the pathogen and this implies that pathogens might take advantage of plant SBT5.2 to degrade their elicitors. The relevance of elicitor degradation is testified by the fact that *Pseudomonas syringae* secretes AprA, a metalloprotease that inactivates flg22, the main elicitor from flagellin (Pel et al., 2014). Selection for destabilization also occurred in Avr4, a protein secreted by the fungal tomato pathogen *Cladosporium fulvum* that is recognised by the Cf-4 immune receptor in tomato. Immune evasion resulted in virulent races that predominantly carry substitutions of Cys residues that destabilise Avr4 in the apoplast, but maintain its ability to protect chitin against chitinases (Joosten et al., 1997; Van den Burg et al., 2003). We were also able to demonstrate that SBT5.2 cleaves the flg22 epitope in flagellin, inactivating its immunogenicity (Buscaill et al., 2024). Interestingly, the fact that both CSP and flagellin are cleaved first in the immunogenic epitopes instead of elsewhere is remarkable and might have resulted from evolutionary pressure to avoid immunity.

Interestingly, we detected some variation in the stability of the csp22 peptides, although all peptides are eventually degraded by SBT5.2. Since the tested peptides all carry the VKWF tetrapeptide before the cleavage site, the differential stability is probably caused by variation in residues after the cleavage site. This indicates that the NDEK tetrapeptide after the cleavage site reduces SBT5.2 sensitivity, whereas the NTSK sequence increases SBT5.2 sensitivity when compared to the NNAK tetrapeptide from CspD. These observations indicate that acidic residues (D or E) at the P2’ and P3’ positions may reduce SBT5.2 sensitivity of csp22 peptides. The different stabilities of the csp22 peptides implies that the relative concentrations of these different csp22 peptides will change after infection and at a distance from the infection site.

A recent pangenomic study on the diversity of csp22 in bacteria revealed that csp22 exhibits a significant copy and epitope variation (Stevens et al., 2023). Interestingly, 25% of the tested csp22 peptides were not immunogenic in tomato carrying the *CORE* receptor and some of these peptides were found to suppress csp22-mediated PTI by blocking the perception of immunogenic csp22. Remarkably, non-immunogenic csp22 peptides have even more significant variation surrounding the S1 site, indicating that antagonistic csp22 peptides might have an enhanced stability to increase their ability to interfere with PTI signaling. These predictions, however, are all based on assays with synthetic csp22 peptides. Although five *CSP* genes are expressed in *Pto*DC3000 during infection, and derived peptides have been detected in crude apoplastic fluids from infected plants by proteomics (Stevens et al., 2023), it remains to be shown which CSPs are perceived during infection. Experiments to delete *CSP* genes or alter the csp22 sequence are challenging because CSPs are collectively essential for bacterial survival (Xia et al., 2001; Graumann et al., 1997) and the RNA binding motif overlaps with the csp22 epitope.

SBT5.2 is the most active subtilase in the apoplast of *N. benthamiana* and its role in degrading csp22 epitopes (this work) and flg22 epitopes (Buscaill et al., 2024) has now been demonstrated. Without preincubation in AF, we did not detect an altered ROS burst upon csp22/flg22 signalling in the *sbt5.2* mutant, indicating that SBT5.2 is not involved in signaling itself. We did also not detect any macroscopic developmental phenotype of *sbt5.2* mutants (Beritza et al., 2024). Likewise, also Arabidopsis *sbt5.2* mutants grow normally (Shardon et al., 2016). Given its abundance and promiscuity, we hypothesize that SBT5.2 might mediate the removal of unstable proteins in the apoplast to maintain extracellular protein homeostasis. The identification of proteins accumulating in the apoplast of *sbt5.2* mutant plants may therefore provide more insights into the endogenous role of SBT5.2.

The fact that a single bacterium produces multiple CSPs that include csp22 peptides that have different stability and immunogenicity indicates that the outcome of interactions involving CSPs is complicated. These interactions will be further complicated by the variation in apoplastic proteases between plant species and upon the secretion of immune proteases. These interactions can be further fine-tuned by the secretion of protease inhibitors by pathogens. Although the csp22 sequences are evolutionary constrained by the fact that they contain the RNA binding motif required for the intrinsic function of CSP (Felix & Boller, 2003), these observations indicate that csp22 variation might underlie a fascinating natural battlefield that is pivotal for the outcome of plant-bacterium interactions.

## Materials & Methods

### Plants

*Nicotiana benthamiana* plants (LAB genotype) were grown at 21°C (night) and 22-23°C (day) under a 16 h light (80-120 μmol/m^2^/s)/ 8 h dark routine in a greenhouse until use.

### Molecular cloning

All used primers and plasmids are summarized in Supplemental **Tables S2 and S2**, respectively. Expression vector pJK155 was generated by cloning inserts of pJP001, pJK120, pFGH029 and pJP002 into pJK082 using a BsaI Golden Gate reaction, resulting in pJK155 (pET28b-T7::OmpA-His-TEV-EPIC1). Open reading frames of *CapB* and *CspD* from *Pto*DC3000 were amplified from genomic DNA of *Pto*DC3000 using primers listed in Supplemental **Table S1** and cloned into pJK155, which was linearized by PCR with primers 5’-gcttggatccggctgctaac-3’ and 5’-accttggaagtataggttttcgtg-3’ using the Gibson ligation method, resulting in pCC03 (pET28b-T7::OmpA-His-TEV-CapB) and pCC04 (pET28b-T7::OmpA-His-TEV-CspD). The open reading frame (ORF) of *Nb*SBT5.2a was commercially synthesized with a C-term 6HIS tag (Twist Bioscience, South San Francisco, California, United States, Supplemental **Table S1**) and assembled with the golden-gate compatible vector pJK001c binary vector (Paulus et al., 2020), 35S promoter module (pICH51288) and 35S terminator module (pICH41414) in a BsaI reaction resulting in expression plasmid pPB097 for transient protein expression of SBT5.2a-His in *N. benthamiana*. This plasmid was transformed *into E. coli* DH10β for amplification, purified and then transformed into *Agrobacterium tumefaciens* GV3101-pMP90. Transformants were selected on plates of LB-agar medium containing 25 µg/ml rifampicin, 10 µg/ml gentamycin and 50 µg/ml kanamycin.

### Expression and purification of CspD

Plasmid pCC04 was transformed into the Rosetta strain of *E. coli* and protein expression was induced with 0.4 mM IPTG at 20 °C and proteins were purified on HisPur ™ Ni-NTA Resin (Thermoscientific) according to the instructions of the manufacturer. Protein purity was verified by protein gel electrophoresis followed by coomassie staining and western blotting using anti-His (HRP) antibody (Miltenyi Biotec). Signals were generated by chemiluminescence using Clarity ECL Western Blotting Substrate (BioRad) and detected with the ImageQuant LAS 4000 (GE Healthcare). Purified proteins were further concentrated using Amicon Ultra centrifugal filter device (3 kDa MW cut-off, Millipore). Protein quantity was measured using Bradford method (Sigma-Aldrich). Proteins were stored in aliquots at -80°C until use.

### Transient protein expression in *N. benthamiana*

To express proteins transiently in *N. benthamiana* by agroinfiltration, overnight cultures of *A. tumefaciens* GV3101(pMP90) carrying binary plasmids to express Epi1 (pFGH048, Grosse-Holz et al., 2018) or SBT5.2a-HIS (pPB097) were harvested by centrifugation. Cells were resuspended in induction buffer (10 mM MgCl2, 10 mM MES pH5.0, and 150 µM acetosyringone) and mixed (1:1) with agrobacteria carrying the silencing inhibitor P19 at OD 600 = 0.5. After 1 h at 21°C, cells were infiltrated with a needleless syringe into the abaxial side of three leaves of 4-week-old *N. benthamiana*. Leaves were harvested and processed at the indicated days after agroinfiltration.

### Isolation of apoplastic fluids

AFs were collected as described previously (Buscaill et al., 2019). Leaves from wild-type, agroinfiltrated or VIGS-silenced plants were submerged in ice-cold water and infiltrated by applying vacuum for 5 min. The surface of water-infiltrated leaves was dried with absorbing paper and leaves were carefully mounted in an empty 20-ml syringe, placed in a 50-ml tube. AFs were collected by centrifugation at 2000 x *g* at 4°C for 20 min and used immediately or flash-frozen and stored at -20°C.

### SBT5.2a-His and P69B-His purification

Four-week-old N.b leaves were infiltrated with a 1:1 mixture (final OD600 = 0.5 for each) of *Agrobacterium tumefaciens* GV3101 containing the silencing suppressor P19 and pPB097 or pJP008. Apoplastic fluid containing SBT5.2a-His or P69B-His was extracted 6 days after infiltration and purified as previously described (Schuster et al., 2022; Homma et al., 2023).

### ROS assays

The ROS burst assay was performed as described (Buscaill et al., 2019) with the difference that L-012 (Wako Chemical, Japan) was used instead of luminol and the diameter of leaf discs used here was 4 mm rather than 6 mm. Briefly, after incubation in water overnight, one leaf disc (4 mm diameter) was added to 100 µl solution containing 25 ng/µl L-012, 25 ng/µl Horse Radish Peroxidase (HRP) and specified elicitor treatments. For assays with elicitors treated with AFs, 500 nM csp22 or purified CspD were incubated in AFs from specified *N. benthamiana* leaves (wild-type, agroinfiltrated or VIGS-silenced) for 1 hour at room temperature with slightly shaking. After incubation, 25 ng/µl L-012 and 25 ng/µl HRP were added to the AFs. Chemiluminescence was measured immediately with the Infinite M200 plate reader (Tecan, Mannedorf, Switzerland) every minute for one hour.

### Peptide synthesis

All used peptides were custom-synthesised by Genscript and are summarized in Supplemental **Table S3**.

### Qcsp8 assays

The quenched Csp octapeptide (Qcsp8, VKWFNNAK) from CspD was commercially synthesized with a DABCYL N-terminal modification and an EDANS C-terminal modification (GenScript, Piscataway, New Jersey, United States) at a purity of 95.7%. It was resuspended in DMSO at a concentration of 1 mM. This stock solution was further diluted in water to a concentration of 200 µM. AFs or purified SBT5.2a or P69B were mixed with Qcsp8 at a final concentration of 10 µM in a volume of 100 µl, and fluorescence was measured immediately or after the indicated incubation time at 21°C using an Infinite M200 plate reader (Tecan, Mannedorf, Switzerland) with an excitation wavelength of 335 nm and emission wavelength of 493 nm.

### *In vitro* degradation assays of CspD by Afs

Purified CspD (stock concentration 10 μM; final concentration 2 μM) were incubated in AFs from specified *N. benthamiana* leaves (wild-type, agroinfiltrated or VIGS-silenced) for indicated time at room temperature with slightly shaking. Proteins were analysed by SDS-PAGE and Coomassie staining.

### Peptide release from digestion of CspD in AF

10 ng/µl of purified CspD protein produced in *E. coli* was incubated in AF from wild-type *N. benthamiana* for 15 min or 60 min at room temperature. CspD and AF alone were used as negative controls and two technical replicates were included for each treatment or control. After incubation, samples for the analysis of endogenously digested peptides in the AF were generated by supplementing the AF with four volumes of MS-grade acetone, followed by incubation on ice for 1 hour and centrifugation at 18,000 x *g* for 15 min. Four-fifths of the supernatants were then transferred to fresh Eppendorf tubes and the acetone evaporated by vacuum centrifugation. The dried peptide samples were then dissolved in 0.1% formic acid and immediately analyzed without further clean-up. LC-MS/MS analysis and peptides identification was performed as described (Buscaill et al., 2019) except that MS/MS spectra data were searched against the sequence of CspD protein. Identified CspD peptides in samples of CspD incubated in AF were normalized by subtraction of the data generated from negative controls and aligned with CspD protein sequence.

### LC-MS/MS

Each sample was analysed on an Orbitrap Elite instrument (Thermo) (Michalski et al. 2012) that was coupled to an EASY-nLC 1000 liquid chromatography (LC) system (Thermo) and an Orbitrap Fusion Lumos (Thermo) coupled to an EASY-nLC 1200 liquid chromatography (LC) system (Thermo). The LC’s were operated in the one-column mode. The analytical column was a fused silica capillary (75 µm × 46 cm) with an integrated fritted emitter (15 µm; CoAnn Technologies) packed in-house with Kinetex C18-XB core shell 1.7 µm resin (Phenomenex). The analytical column was encased by a column oven (Sonation) and attached to a nanospray flex ion source (Thermo). The column oven temperature was adjusted to 50 °C during data acquisition. The LC was equipped with two mobile phases: solvent A (0.2% formic acid, FA, in water) and solvent B (0.2% FA, 19.8% water and 80% acetonitrile, ACN). All solvents were of UPLC grade (Honeywell). Peptides were directly loaded onto the analytical column with a maximum flow rate that would not exceed the set pressure limit of 980 bar (usually around 0.6 – 1.0 µL/min). Peptides were subsequently separated on the analytical column by running a gradient of solvent A and solvent B. The exact composition of the gradient and the settings for the mass spectrometers can be found in **Supplemental Materials**.

### Peptide and Protein Identification using MaxQuant

RAW spectra were submitted to an Andromeda (Cox et al., 2011) search in MaxQuant (2.0.2.0.) using the default settings (Cox & Mann 2008). Label-free quantification and match-between-runs was activated (Cox et al., 2014). The MS/MS spectra data were searched against a project specific database containing 2 sequences of interest (ACE_0686_SOI_v01.fasta; 2 entries). All searches included a contaminants database search (as implemented in MaxQuant, 245 entries). The contaminants database contains known MS contaminants and was included to estimate the level of contamination. Andromeda searches allowed oxidation of methionine residues (16 Da) and acetylation of the protein N-terminus (42 Da). No dynamic modifications were selected. Enzyme specificity was set to “unspecific”. The instrument type in Andromeda searches was set to Orbitrap and the precursor mass tolerance was set to ±20 ppm (first search) and ±4.5 ppm (main search). The MS/MS match tolerance was set to ±0.5 Da. The peptide spectrum match FDR and the protein FDR were set to 0.01 (based on target-decoy approach). Minimum peptide length was 6 amino acids and maximum amino acid length 36. For protein quantification unique and razor peptides were allowed. Modified peptides were allowed for quantification. The minimum score for modified peptides was 40. Label-free protein quantification was switched on, and unique and razor peptides were considered for quantification with a minimum ratio count of 2. Retention times were recalibrated based on the built-in nonlinear time-rescaling algorithm. MS/MS identifications were transferred between LC-MS/MS runs with the “match between runs” option in which the maximal match time window was set to 0.7 min and the alignment time window set to 20 min. The quantification is based on the “value at maximum” of the extracted ion current. At least two quantitation events were required for a quantifiable protein. Further analysis and filtering of the results was done in Perseus v1.6.10.0. (Tyanova et al., 2016). Comparison of protein group quantities (relative quantification) between different MS runs is based solely on the LFQ’s as calculated by MaxQuant, MaxLFQ algorithm (Cox et al., 2014).

### Virus-induced gene silencing (VIGS)

*N. benthamiana* plants were silenced for *NbCORE*, *SBT5.2*, glucuronidase *GUS* (negative control) and phytoene desaturase *PDS* (positive control) were generated using VIGS as previously described (Dodds et al., 2023, Beritza et al., 2024). Briefly, overnight cultures of *A. tumefaciens* GV3101 were collected and resuspended in agroinfiltration buffer (10 mM MgCl2, 10 mM MES pH 5.0, and 100 mM acetosyringone). Suspensions of bacteria containing *TRV2gg::NbCORE*, *TRV2::SBT5.2* and *TRV2::GUS* were mixed 1:1 separately with bacteria containing *TRV1* at OD600 = 0.5 for each bacteria. After incubation for 1 hour at room temperature, the mixed cultures were infiltrated into true leaves of 2 week-old *N. benthamiana* plants. The infiltrated seedlings were grown in a growth chamber until use.

### Labelling of active Subtilases

FP-TAMRA (Thermo Scientific) was prepared as 10 µM stock solutions in dimethyl sulfoxide. Labelling was performed as described previously (Buscaill et al., 2019). For fluorescence gel imaging, the apoplastic fluids were incubated with 0.2 µM probes for 1 h at room temperature in the dark. The labelling reactions were stopped by adding 4x gel loading buffer (200 mM Tris-HCl (pH6.8), 400 mM DTT, 8% SDS, 0.4% bromophenol blue, 40% glycerol) and heating at 90°C for 5 min. The proteins were separated on SDS-PAGE and fluorescence was detected from protein gels using the Typhoon FLA 9000 scanner (GE Healthcare Life Sciences) using Cy3 settings (532nm excitation and 610PB filter).

### Infection assays

csp22 peptides were diluted in water. Three fully expanded leaves of 3-4 weeks old *N. benthamiana* plants were infiltrated with the different concentrations of csp22 peptide or with water as a mock control. 24h later, infiltrated leaves were infiltrated with 10^5^ CFU/ml *Pto*DC3000(ΔhQ) or *Pta*6605. The next day (1 dpi), three leaf discs were punched with a cork borer from each infected leaf, and surface-sterilised with 15% hydrogen peroxide for 2 minutes. Leaf discs were then washed twice in MilliQ and dried under sterile condition. Leaf discs were placed into a 1.5 ml safe-lock Eppendorf tube with three 3 mm diameter metal beads and 1 ml of MilliQ. Tubes were placed in tissue lyser for five minutes at 30 Hertz/second. 20 µl of undiluted tissue and serial dilutions were plated on LB-agar plates containing Pseudomonas CFC Agar Supplement (Oxoid SR0103). Plates were allowed to dry, incubated at 28°C for two days and then colonies were counted. The *p*-value was calculated using the two-tailed Student *t*-test to compare bacterial growth between leaves from WT and *sbt5.2s* mutant plants.

### Statistics

All values shown are mean values, and the error intervals shown represent standard error of the mean (SE), unless otherwise indicated. All experiments have been reproduced, and representative data sets are shown.

### Phylogenetic analysis

Proteomes of *Pseudomonas syringae* strains were obtained from the pseudomonas.com database (Winsor et al., 2016) and proteomes from *Ralstonia solanacearum* and *Xanthomonas campestris* were obtained from RefSeq database (O’Leary et al., 2016). Cold shock domain-containing proteins were identified using hmmsearch function from HMMER 3.3.2 (Finn et al., 2011) with PF00313 (cold-shock DNA-binding domain) Pfam profile (Mistry et al., 2021). Only sequences with recognisable csp22 epitope were retained. Protein sequences were aligned using MAFFT 7 with L-INS-i algorithm (Katoh & Standley, 2013). A maximum likelihood phylogenetic tree was constructed using IQ-TREE 2 (Minh et al., 2020) with the best-fit substitution model from ModelFinder (Kalyaanamoorthy, et al., 2017). Branch support values were calculated using ultrafast bootstrap with 1000 replications. Phylogenetic trees were visualised using iTOL (Lurenic & Bork, 2021) with midpoint rooting.

## Acknowledgements

We like to thank Jiorgos Kourelis for cloning pJK155; Urszula Pyzio for excellent plant care; and Sarah Rodgers, Patricia Bowman and Caroline O’Brien for excellent technical support.

## Author contributions

CC performed most of the experiments; PB co-supervised the project and performed infection assays and Qcsp8 experiments; FK and MK generated proteomics data; JH purified SBT5.2 and performed the CspD cleavage assay; NS performed ROS assays presented in Figures 7 and 8; RH conceived and supervised the project; RH and CC wrote the article with input from all co-authors.

## Data availability

The mass spectrometry proteomics data for the on-bead digestions have been deposited to the ProteomeXchange Consortium via the PRIDE (Vizcaíno et al. 2016) partner repository (https://www.ebi.ac.uk/pride/archive/) with the dataset identifier PXD048912. During the review process the data can be accessed via a reviewer account (Username; Password: ErLKUZ3o)

---

**Figure S1.**
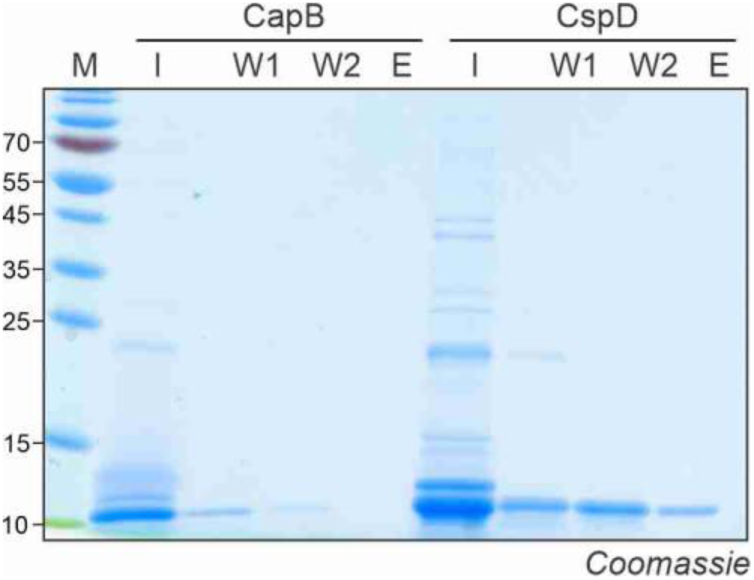
Expression and purification of CapB and CspD. CapB and CspD with expressed with N-terminal His-TEV peptides in *E. coli* and purified over Ni-NTA. I, input sample; W1, wash with 100 mM imidazole; W2, wash with 150 mM imidazole; E, eluate with 250 mM imidazole. Samples were separated by SDS-PAGE and stained with Coomassie.

**Figure S2.**
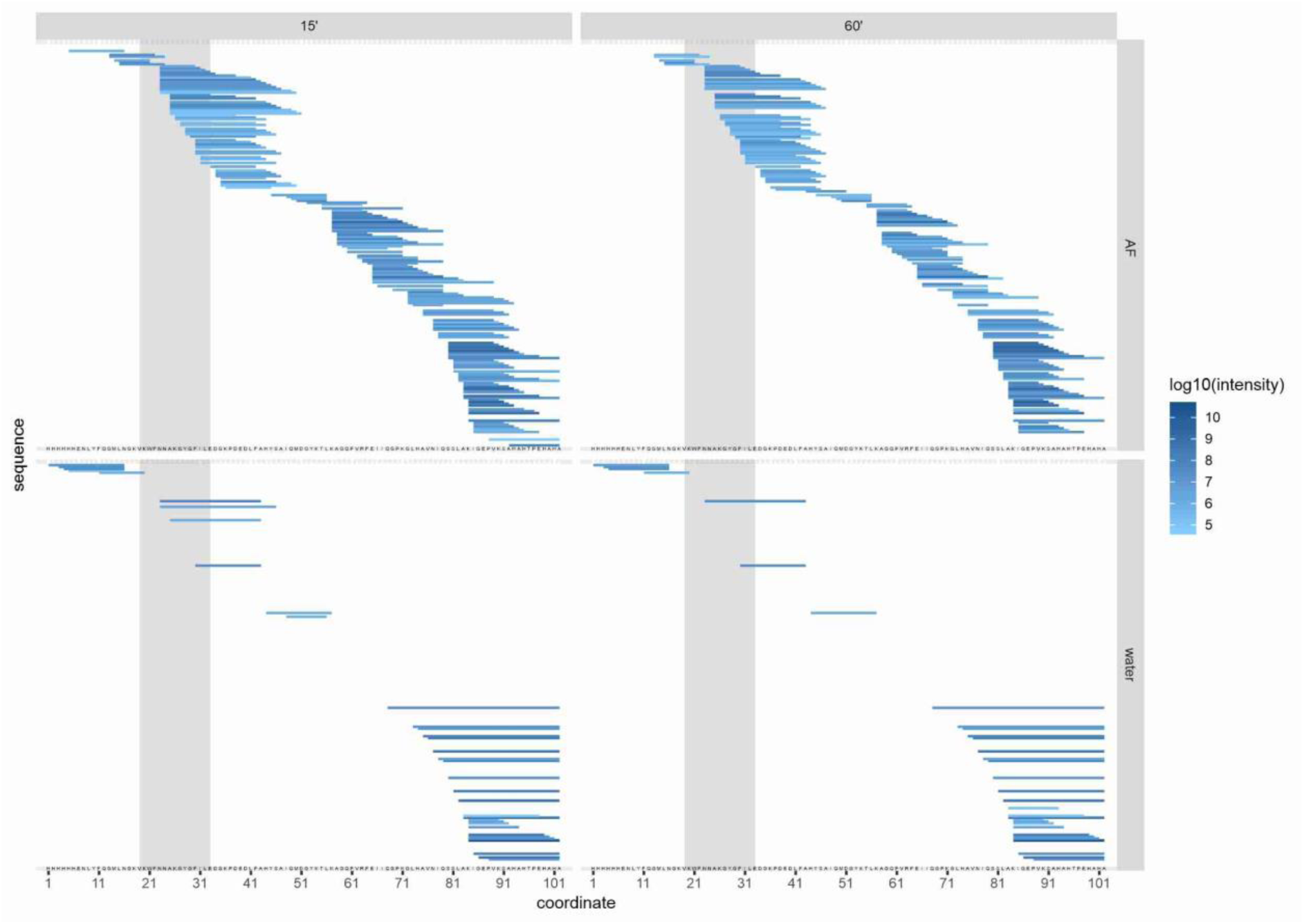
Detected CspD peptides. Coverages of detected CspD peptides after incubation of CspD in water (bottom) or AF (top) for 15 (left) and 60 (right) minutes.

**Figure S3.**
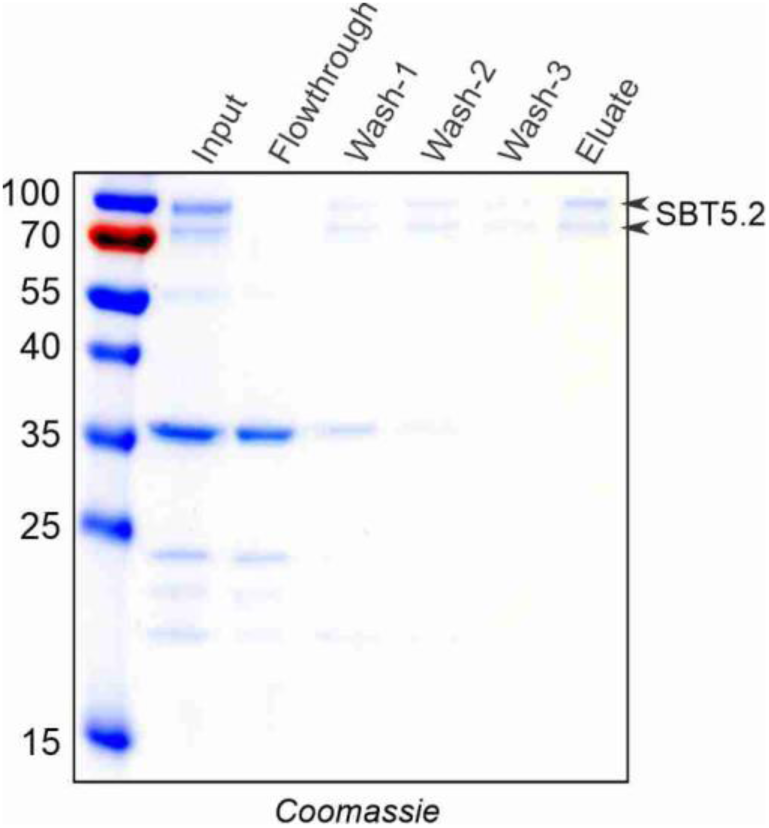
Purification of SBT5.2a-His from AF of agroinfiltrated plants. SBT5.2a-His was transiently expressed by agroinfiltration of *N. benthamiana*. AF was isolated at 6 dpi and purified over Ni-NTA. The column was washed with 50 mM imidazole and eluted with 250 mM imidazole.

**Figure S4.**
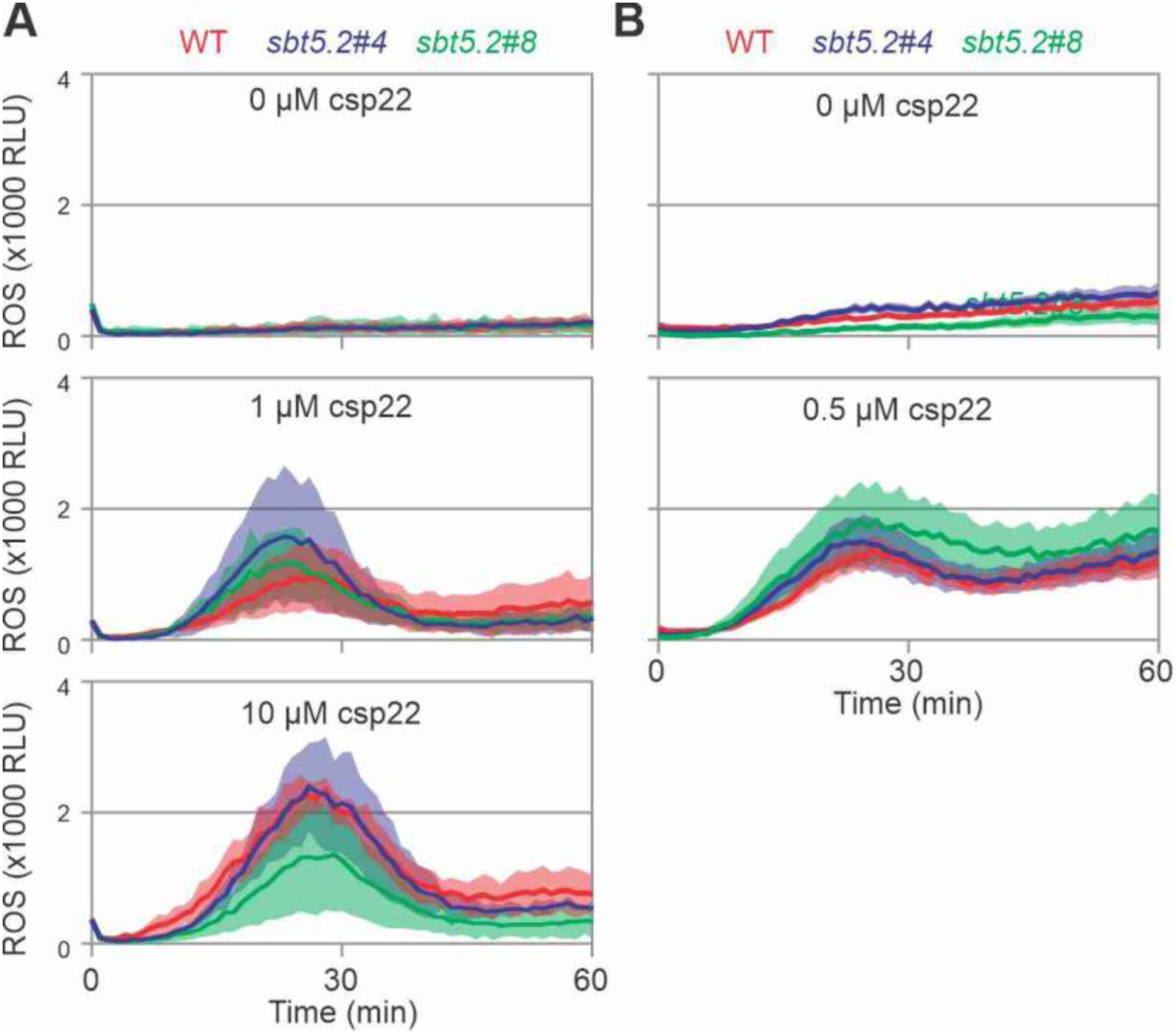
Mutant *sbt5.2* plants still sense csp22. **(A)** Leaf discs of 7-week old plants were treated with 0, 1 or 10 μM csp22 and luminescence with luminol was measured for one hour in relative light units (RLU). Error shades represent the STDEV of n=4 replicates. **(B)** Leaf discs of 7-week old plants were treated with 0, 1 or 10 μM csp22 and luminescence with LO-12 was measured for one hour in relative light units (RLU). Error shades represent the SE of n=6 replicates.

**Figure S5.**
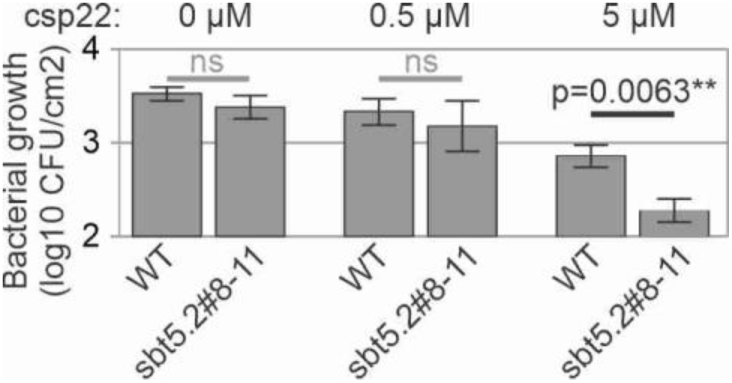
Immune priming is increased in the *sbt5.2* mutant. Leaves of WT and *sbt5.2* mutant *N. benthamiana* were infiltrated with water or 0.5 or 5 μM csp22 and incubated for 24 hours and then injected with 1x10^5^ bacteria/mL of the flagellin mutant strain of *Pseudomonas syringae* pv. *tabaci* 6605 (*Pta*6605*ΔfliC*). Colony forming units (CFUs) were determined one day post infection (dpi). Error bars represent SE of n=6 replicates.

**Figure S6.**
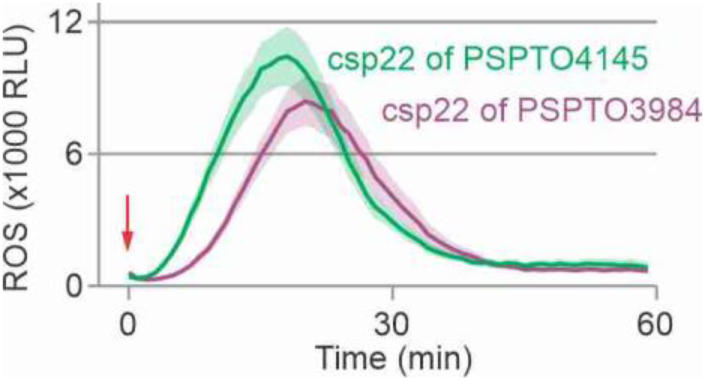
Also csp22 peptides from other CSPs trigger an oxidative burst. Leaf discs of 6-week old plants were treated with 500 nM csp22 of PSPTO3984 or PSPTO4145, and ROS was measured in relative light units (RLU). Error shades represent the SE of n=6 replicates.

**Figure S7.**
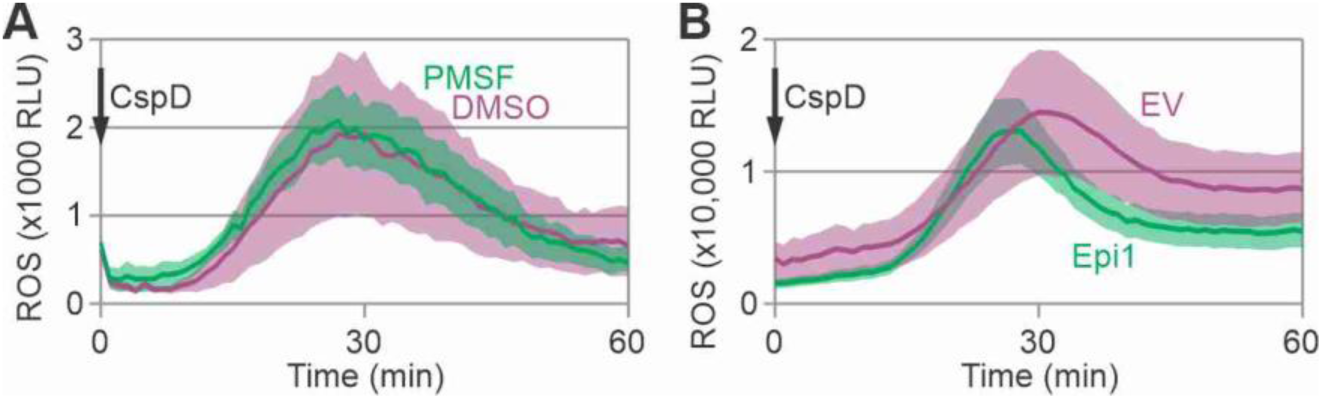
Subtilase inhibitors do not block CspD perception. **(A)** Leaf discs of *N. benthamiana* were treated with 0.5 mM PMSF or 1% DMSO when floating on luminol-HRP solution and were triggered with 500 nM CspD protein and the release of reactive oxygen species (ROS) was detected by luminescence. Error shades represent SE of n=6 replicates. **(B)** Leaf discs taken from agroinfiltrated leaves expressing Epi1 or empty vector (EV) control, taken at 3 dpi, floating on luminol-HRP solution, were triggered with 500 nM CspD protein and the release of reactive oxygen species (ROS) was detected by luminescence. Error shades represent SE of n=6 replicates.

**Table S1.**
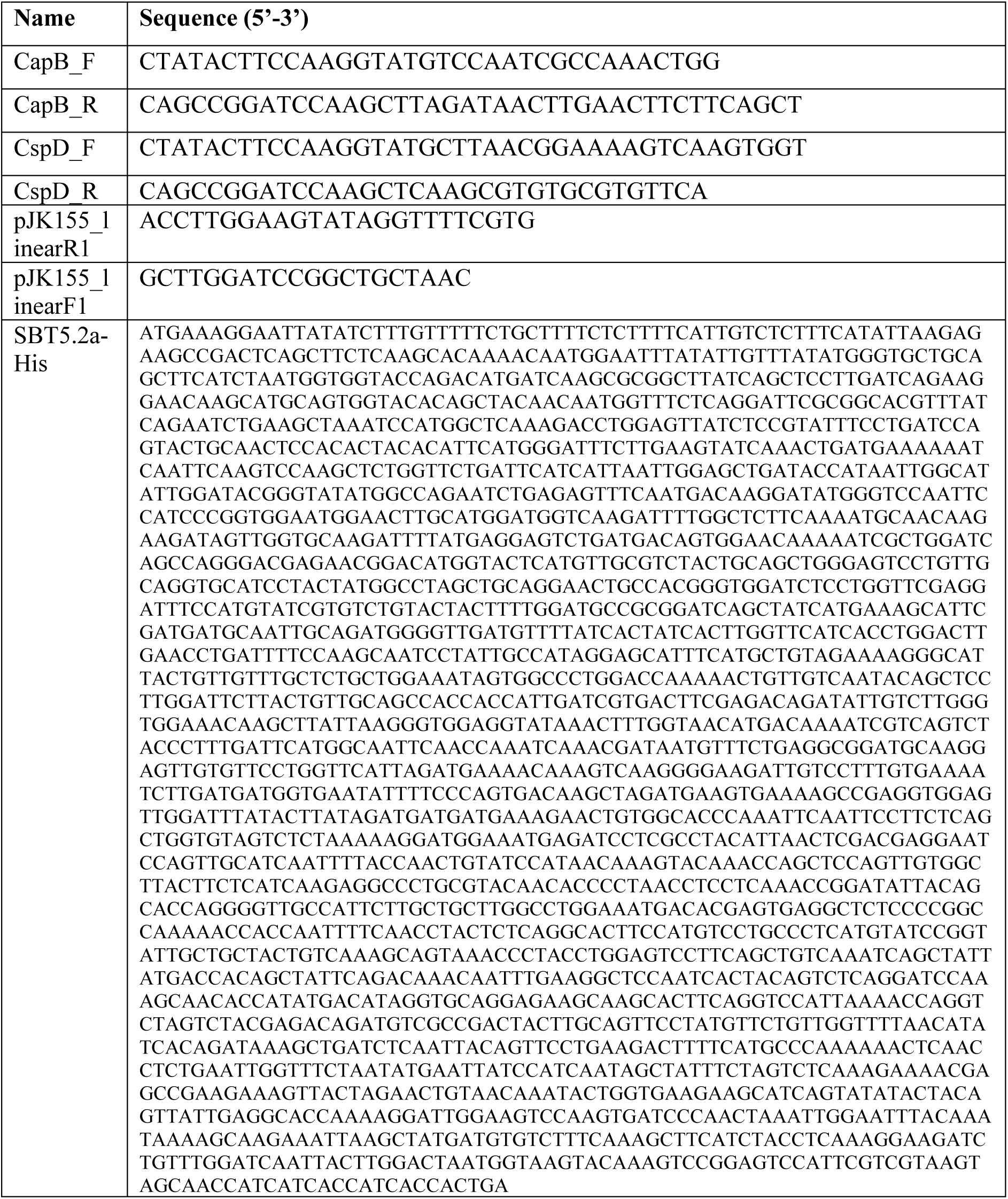
Used synthetic oligonucleotides.

**Table S2.**
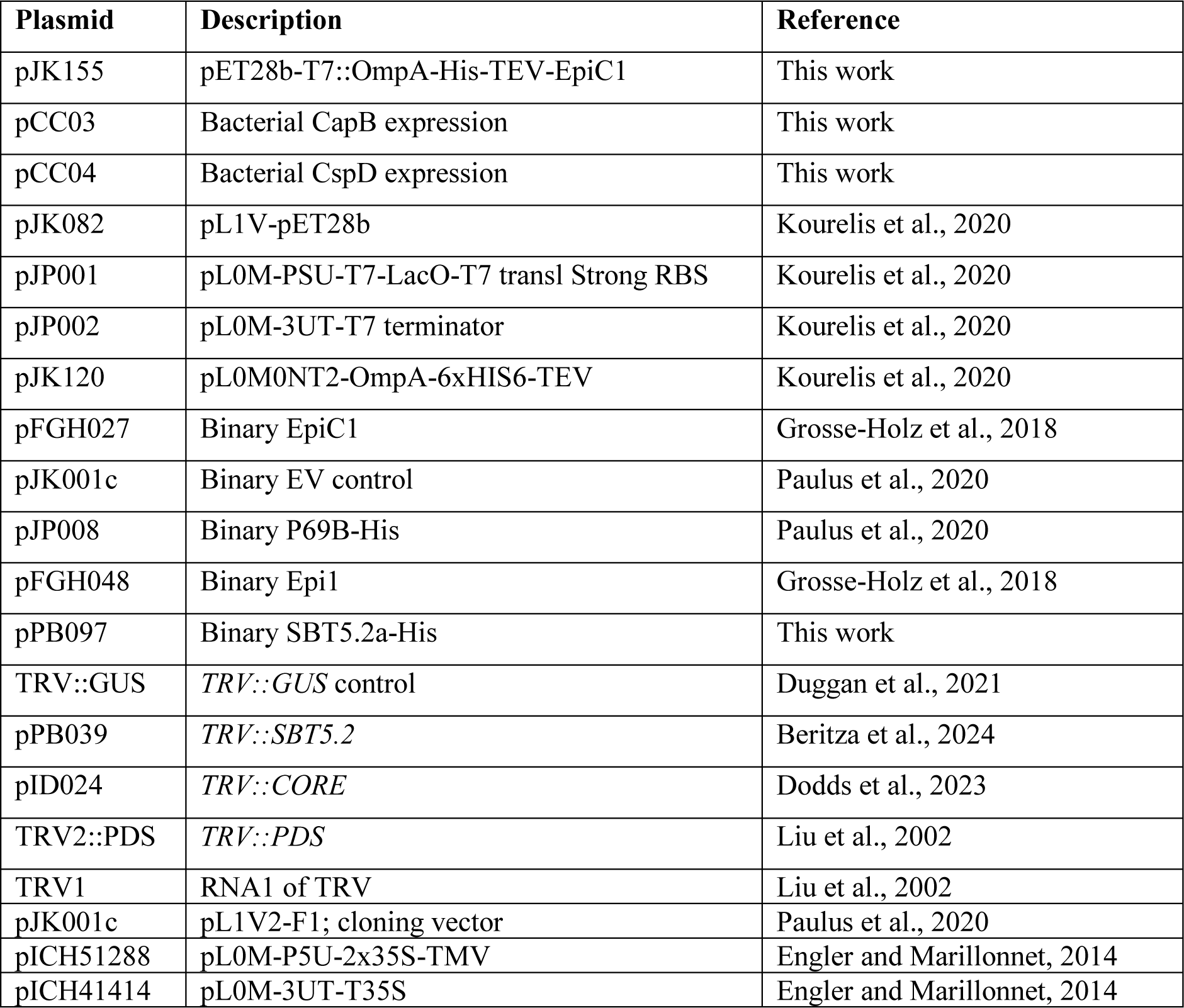
Used plasmids.

**Supplemental Table S3.** Used synthetic peptides.

| Peptide | Description | Manufacturer |
| --- | --- | --- |
| LNGKVKWFNNAKGYGFILEDGK | csp22 (CspD) | Genscript |
| DabcyI-VKWFNNAK-Edans | Qcsp8 | Genscript |
| DTGTVKWFNTSKGFGFISRDSG | csp22 PSPTO3984 | Genscript |
| QTGTVKWFNDEKGFGFITPQSG | csp22 PSPTO4145 | Genscript |

